# Perceptual Resolution of Ambiguity: Can Tuned, Divisive Normalization Account for both Interocular Similarity Grouping and Difference Enhancement

**DOI:** 10.1101/2024.04.01.587646

**Authors:** Jaelyn R. Peiso, Stephanie E. Palmer, Steven K. Shevell

## Abstract

Our visual system usually provides a unique and functional representation of the external world. At times, however, the visual system has more than one compelling interpretation of the same retinal stimulus; in this case, neural populations compete for perceptual dominance to resolve ambiguity. Spatial and temporal context can guide perceptual experience. Recent evidence shows that ambiguous retinal stimuli are sometimes resolved by enhancing either similarity or differences among multiple percepts. Divisive normalization is a canonical neural computation that enables context-dependent sensory processing by attenuating a neuron’s response by other neurons. Experiments here show that divisive normalization can account for perceptual representations of either similarity enhancement (so-called grouping) or difference enhancement, offering a unified framework for opposite perceptual outcomes.

## Introduction

Visual neural processing achieves functional percepts amidst the ambiguity inherent in under-constrained sensory signals (Brascamp & Shevell, 2021). For example, consider how we effortlessly navigate a three-dimensional (3-D) world, complete with depth, 3-D obstacles, and moving objects. We achieve this complex task despite our retinae receiving only two-dimensional (2-D) projections of the external world. This projection of the 3-D world into 2-D retinal signals, and subsequently back to 3-D perceptual representations, inherently results in information loss. Consequently, multiple interpretations can arise for the external stimulus based on its retinal representation. The visual system maintains perceptual continuity in typical viewing conditions by leveraging local context and configural cues and employing selective attention. For example, local context constrains perception by adapting the sensory signal to align with the local statistics in the image (Coen-Cagli, Kohn, & Schwartz, 2015; Lee & Mumford, 2003). Configural cues anchor perception in the learned statistical patterns of the external environment (Fowlkes, Martin, & Malik, 2007; Lee & Mumford, 2003). Spatial attention offers additional constraints by selectively amplifying sensory signals from regions of focus (Desimone & Duncan, 1995; Hillyard, Vogel, & Luck, 1998; Ling & Blake, 2012). Under-constrained sensory signals can result in multi-stable perception without these supplementary constraints, as observed in classic visual ‘illusions’ (Murray & Adams, 2019) or the perceptual resolution of binocularly rivalrous stimuli.

In binocular rivalry, incompatible stimuli are presented to corresponding retinotopic regions of the two eyes. Such interocular conflict often induces perceptual alternations like those observed in other forms of bistable perception (Brascamp, Qian, Hambrick, & Becker, 2019; Cao, Wang, Sun, Engel, & He, 2018). Intriguingly, when two spatially separated rivalrous patches are present, they often resolve as identical during perceptual alternations, a phenomenon referred to as interocular grouping (Logothetis et al., 1996). Interocular grouping demonstrates a flexible, similarity-enhancing bias. It occurs with multiple, steadily presented rivalrous stimuli and even when the rivalrous images are swapped between the two eyes several times each second. The prevalence of this perceptual similarity bias has prompted theories proposing a grouping process driven by a posited mechanism for spatially integrated ‘filling-in’ that is regulated by attention (Brascamp et al., 2019; Buckthought, Kirsch, Fesi, & Mendola, 2021; Lee & Blake, 1999; Lee & Blake, 2004). Emerging evidence, however, adds nuance to this view. Perceptual resolution, in some circumstances, can be biased toward resolving the multiple regions such that they appear as dissimilar as possible (that is, difference enhancement). The experiments reported here test whether a single theoretical framework, chromatically-tuned divisive normalization, can account for both similarity and difference enhancement, depending on context.

Divisive normalization is well-documented in sensory neurons and appears as a context-dependent nonlinearity in many brain areas (Coen-Cagli, Kohn, & Schwartz, 2015; Louie & Glimcher, 2019; Schwartz & Simoncelli, 2001). It is widely considered a foundational computation for information processing within the brain (Carandini & Heeger, 2011; Louie & Glimcher, 2019; Olsen, Bhandawat, & Wilson, 2010; Schwartz & Simoncelli, 2000). In its simplest form, divisive normalization means applying a gain factor to a cell’s response with the value of the gain set by other nearby neural responses (Carandini & Heeger, 2012; Heeger, 1992; Carandini, Heeger, & Movshon, 1997; Schwartz & Simoncelli, 2001). Divisive normalization provides a mechanism for redundancy reduction, so it is consonant with the efficient coding hypothesis (Barlow, 1961). Divisive normalization also provides a biologically plausible mechanism to explain non-linear neural responses, such as surround suppression (Simoncelli & Schwartz, 1999) and cross-orientation suppression (Brouwer & Heeger, 2011). In both cases, the outputs of classical receptive-field signals are attenuated by the neural response to the surround or by the neural response to the superposition of a non-preferred stimulus. Similarly, divisive normalization has been suggested to underlie the mutual inhibition used to explain the temporal dynamics of binocular rivalry (Freeman, 2005; Lehky, 1988; Ling & Blake, 2012; Wilson, 2003). In these models, neurons responding to each eye’s stimulus inhibit each other through a shared normalization pool (Ling & Blake, 2012; Said & Heeger, 2013). Specifically, one representation dominates perception while the normalization term suppresses the other representation.

Models of binocular rivalry focus on rivalry dynamics, using noise (e.g., Brascamp et al., 2006; Freeman, 2005; Kim et al., 2006; Moreno-Bote et al., 2007) or the combination of noise and adaptation (e.g., Lehky, 1988; Lankheet, 2006; van Ee, 2009; Wilson, 2001; Wilson, 2003) to model the dynamic perceptual alternations characteristic of rivalry. Some models posit a fundamental role for attention in binocular rivalry oscillatory dynamics, either through gating visual awareness or as a model extension to account for interocular grouping (Brascamp & Blake, 2012; Dieter, Brascamp, Tadin, & Blake, 2016; Hancock & Andrews, 2007; Lee & Blake, 2004; Li et al., 2017; Ling & Blake, 2012; Said & Heeger, 2013). Some normalization models of binocular rivalry posit a shared pool of neurons conveying information from different eyes with dissimilar feature-tuning. These models are consistent with a ubiquitous grouping process, as the representations of common signals will always dominate less pervasive signals when the two mutually impose divisive strength (Lee & Blake, 1999). While existing models can account for interocular grouping, they cannot account for maximally differentiated percepts that dominate perception despite the physical possibility of forming grouped (similarity-enhanced) percepts (Peiso & Shevell, 2020).

The development of the framework here is rooted in previous findings showing that similarity enhancement can be strongly attenuated, and perceptual difference enhancement increased even with minor changes to classic stimuli used to demonstrate similarity enhancement (Peiso & Shevell, 2020). These findings provide initial evidence against a ubiquitous similarity grouping process operating on ambiguous neural representations. This model diverges from some existing theories by not relying on top-down mechanisms, such as selective attention, to explain interocular grouping (Lee & Blake, 2004; Said & Heeger, 2013). Instead, it employs a simple, stimulus-driven gain change underpinned by feature-tuned divisive normalization. Feature-tuned divisive normalization is a process whereby a neuron’s response is modulated by neurons with similar response properties (e.g., orientation, color, motion) rather than neurons that are spatial neighbors (Bloem & Ling, 2019; Klímová, Bloem, & Ling, 2023).

To test the model with feature-tuned normalization, experiments varied a chromatic background while keeping rivalrous regions constant. The assumption is that neurons with similar tuning are pooled across the visual field and so influence perceptual resolution during binocular rivalry, regardless of their possible interocular conflict. Stimuli resembled those that traditionally induce interocular similarity grouping, with each of the two separated rivalrous regions containing the same chromatic rivalry (e.g., Slezak & Shevell, 2018; Peiso & Shevell, 2020). In various conditions the background could be chromatically congruent with one of the rivalrous signals, chromatically neutral, or chromatically rivalrous itself. The experiments test whether a change in background context influences the perceptual resolution of ambiguity, shifting it from similarity enhancement to difference enhancement, in line with predictions from the divisive normalization framework. The experiments also test whether such changes in perceptual outcomes can be attributed to the chromatic contrast between the rivalrous regions and their background.

## General Methods

### Apparatus

Stimuli were displayed on a calibrated NEC MultiSync FP2141SB cathode ray tube (CRT) monitor driven by an iMac computer. Observers viewed the CRT through an eight-mirror haploscope, which presented different stimuli to corresponding retinotopic regions in each eye (Figure 1). A chin rest was employed to maintain an approximately 115-cm-long light path through the haploscope. To ensure stable fusion of the two images and account for individual differences in interocular distance, observers adjusted the position of the final mirror set. Two Nonius lines facilitated image fusion, with the left eye presented with top and left Nonius lines and the right eye presented with bottom and right Nonius lines. A properly fused image exhibited one fixation point, horizontally aligned left and right Nonius lines, vertically aligned top and bottom Nonius lines, and a binocularly fused square frame.

**Figure 1:**
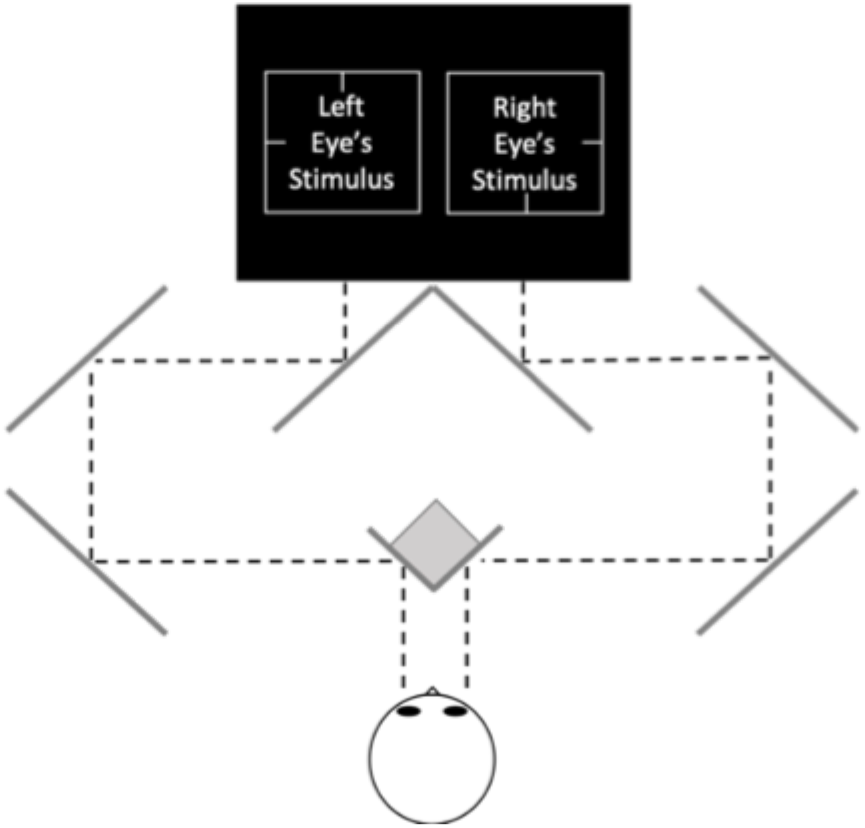
Mirror haploscope display. The CRT display is represented by a black rectangle. The eight mirrors are depicted by

### Observers

Five observers (three female; age 23–32) provided written informed consent prior to participation, as required by the University of Chicago’s Institutional Review Board. Observers were screened for normal stereoscopic vision using the Titmus Stereo Test and for normal color vision employing Ishihara Plates and Rayleigh matches made with a Neitz anomaloscope. Data for both experiments were collected concurrently. All observers participated in both experiments. All observers were naïve to the experimental hypotheses.

### Stimuli

Individualized isoluminant stimuli were generated for each observer to minimize luminance differences. To achieve this, observers repeatedly performed heterochromatic flicker photometry (HFP), a method used to measure the spectral sensitivity of the human eye and define the human photopic luminosity function (Bone & Landrum, 2004; Lee, Martin, & Valberg, 1988; Wyszecki & Stiles, 1982). The HFP stimulus involved a single region (i.e., disk) oscillating at approximately 15 Hz between two distinct chromaticities. Observers adjusted the intensity of one light to minimize the perception of flicker.

Each observer completed five HFP repetitions for three chromaticity pairs: red-appearing/green-appearing (R/G), blue-appearing/green-appearing (B/G), and blue-appearing/red-appearing (B/R) on three separate days. The five measurements per color pair were averaged, resulting in three daily means. After taking the average of these daily means, a final analytical check was made by using the R/G and B/G equiluminant ratios to calculate a predicted B/R ratio. This calculated B/R ratio was compared to the measured B/R ratio, allowing for a deviance of ±10%.

All stimuli were generated in MATLAB® as indexed images for efficient rendering. Chromatically defined stimuli were presented in Interocular-Switch Rivalry (ISR), which entails swapping stimuli between the two eyes at a frequency of 3.75 Hz, or 7.5 swaps per second (Christiansen, D’Antona, & Shevell, 2017; Logothetis, Leopold, & Sheinberg, 1996). Non-rivalrous regions, such as the fixation point and Nonius lines, remained constant across all trials. Based on previous studies that found no significant differences in dominance times between conventional (Figure 2A) and patchwork (Figure 2B) stimulus configurations when presented in ISR (Slezak & Shevell, 2018; Peiso & Shevell, 2020; Lange & Shevell, 2020), only patchwork stimuli requiring interocular integration were tested (Figure 2B). A conventional presentation classically refers to when each eye receives similar signals for each rivalrous region (Figure 2A). A patchwork presentation refers to when each eye receives a different signal for each rivalrous region (Figure 2B).

**Figure 2:**
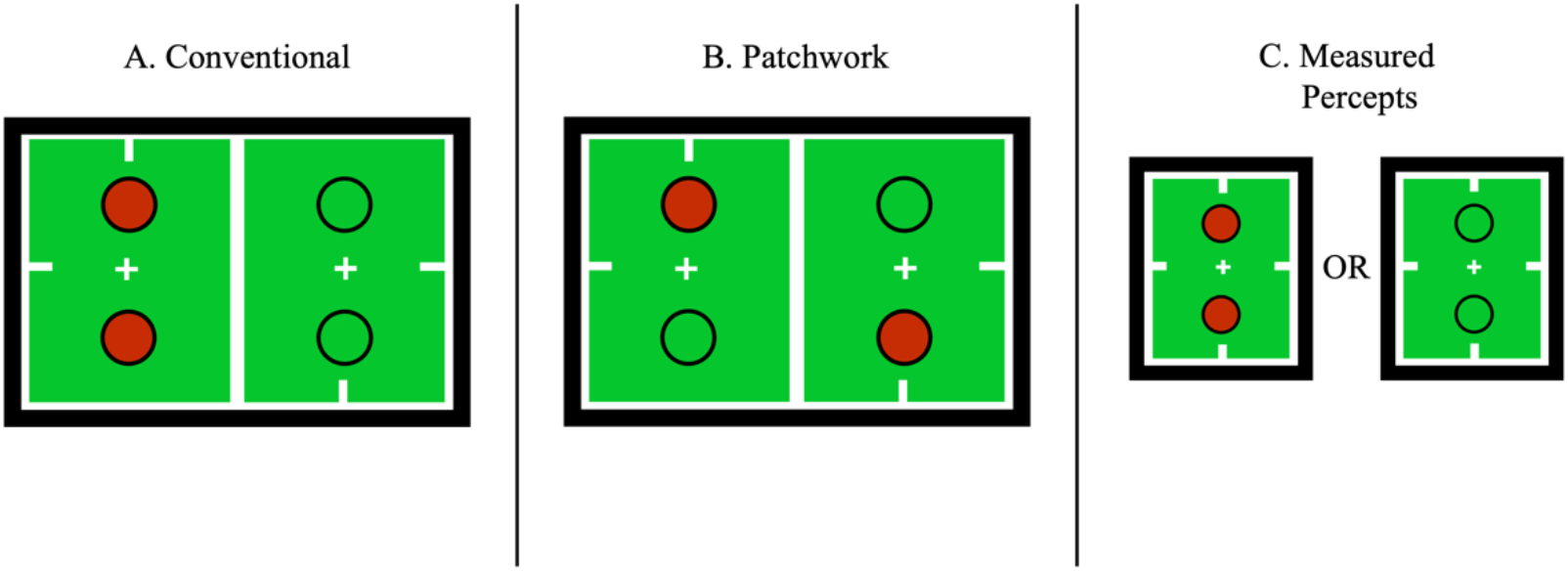
Conventional and patchwork presentations A) Conventional presentation refers to a stimulus with each eye receiving an identical signal for rivalrous regions. B) Patchwork presentation refers to a stimulus with each eye receiving a different signal for rivalrous regions. C) Measured percepts from stimuli A and B.

All stimulus arrays shared the same arrangement while chromaticity and rivalry status were varied. Each array featured one rivalrous region 1.5° above fixation and one 1.5° below fixation (Figure 2), displayed within a 1.5° aperture. All stimuli were presented inside 4.5° by 4.5° fusion boxes with Nonius lines. Fusion box edges were at a chromaticity metameric to the equal-energy spectrum and had luminance contrast (Y = ∼25 cd/m^2^) relative to their interior background (called “white”). The chromaticities for all conditions were set at [L/(L + M), S/(L + M)] values of [0.62, 0.3], referred to as “green,” and [0.71, 0.3], referred to as “red” (MacLeod and Boynton, 1979). The “grey” value was [0.665, 1.0]. The unit of [S/(L + M)], which is arbitrary, was set to 1.0 for equal-energy-spectrum “white.” Red, green, and gray-appearing chromaticities were set at approximately 15 cd/m^2^. All stimuli had two vertically oriented rivalrous regions with dark annuli (Y = ∼0.1 cd/m^2^), increasing the total visual angle of disks with annuli to 1.75°. Annuli were included to aid fusion and to separate background and disk rivalrous regions in Experiment 2.

### Experimental Protocol

The experimental protocol was identical for both experiments. The trial order was randomized for each observer on each experimental day. At the onset of each experimental day and after any scheduled or requested breaks, observers dark-adapted by sitting in the experiment suite for 5 minutes prior to data collection, with all light-emitting electronics turned off. Instructions were displayed on the screen using images to indicate target percepts. Text instructions indicated the gamepad buttons corresponding to each target percept. During each trial, observers were instructed to press and hold buttons on a gamepad for the duration that they experienced each measured percept (described below). Observers were instructed to withhold button presses for all percepts not indicated by the instructions, including partially resolved or piecemeal percepts.

Total dominance durations were calculated by taking the average dominance duration of each measured percept for each of the three days. Standard errors of the mean were calculated using the mean total dominance durations for each of the three experimental days. Each trial began with an instruction screen that indicated which button to press for each measured percept. To reduce the possible impact of differential adaptation between the two eyes from the onset ISR phase and potential onset effects (Carter & Cavanagh, 2007), measurements began after the initial 10 seconds and continued for 60 seconds. The first session was always a practice session, and these data were not analyzed. Practice was followed by three experimental sessions, during which observers completed the same trials in a different random order each day.

### Experiment 1

Peiso and Shevell (2020) unexpectedly found diminished similarity-enhancement (grouping) combined with enhanced difference-enhancement. Can this be explained by pooled divisive normalization? Experiment 1 aimed to determine if non-rivalrous, chromatically stable background signals would pool with like signal components from rivalrous regions and influence perceptual resolution. The normalization framework predicts a reweighting of the visual signal such that less pervasive signals have an increased likelihood of dominating perception during rivalry. In other words, divisive normalization enhances differences, so a disk in red/green rivalry presented on a stable green background is more likely to resolve as red perceptually. The possibility that any observed difference is merely due to chromatic contrast and unrelated to chromatically-tuned normalization is addressed by comparisons with a grey-appearing background. Divisive normalization predicts measurements that resemble a staircase because normalization strength increases with the spatial prevalence of a specific chromatic signal. In the case of a red background, for example, neural populations responding to the background are pooled with the neurons responding to the red component of the rivalrous signal, imposing stronger attenuation on the pool and decreasing the likelihood that rivalrous regions will be perceived as red.

### Stimuli and Procedure

Experiment 1 featured stimuli with non-rivalrous, dichoptically stable backgrounds (Figure 3 A-C). Three experimental conditions were included (Figure 3 A-C), each with two types of measured percepts: 1) both disks resolved as green, or 2) both disks resolved as red. For conditions with chromatic backgrounds (Figure 3 A-B), disks resolved as the same color as their background were considered similarity-enhanced (Figure 3 E & G), and disks resolved as a contrasting color to their background were considered difference-enhanced (Figure 3 D & F). Stimulus C had a neutral grey-appearing background (Figure 3 C) with the same red/green rivalrous disks. Measured percepts (Figure 3 H & I) for stimulus C were considered a baseline for resolving rivalrous disks as red or green.

**Figure 3:**
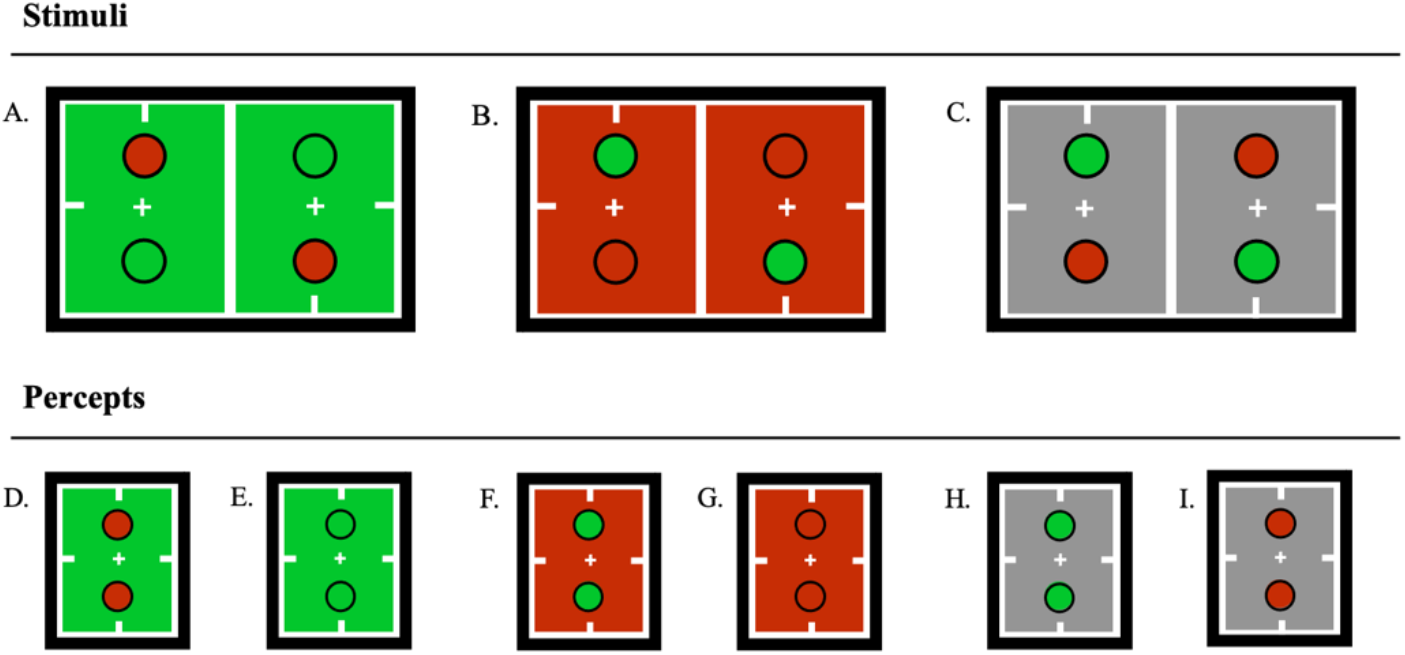
Stimuli and measured percepts for Experiment 1. All stimuli were presented in patchwork. (A): Rivalrous disks within a stable green-appearing context. (B): Rivalrous disks within a stable red-appearing context. (D & F): Measured difference-enhanced percepts for stimuli A and B. (E & G): Measured similarity-enhanced percepts for stimuli A and B. (C): Depicts rivalrous disks within a stable grey-appearing background. (H & I): Measured percepts for stimuli C.

## Results

Experiment 1 was designed to test a very specific prediction of relative dominance durations for red/green rivalrous disks. Specifically, disks should resolve as color-contrasted against their background the most and resolve as the same color as the background the least. Grey backgrounds were expected to elicit intermediate dominance durations, serving as a baseline for comparison. When grouped by resolved disk color, each observer’s data should resemble two staircases. To capture the relational context of the prediction, a non-parametric group analysis used a one-tailed binomial test to assess the binomial probability of the observed results by chance under the null hypothesis (H_0_) of a random order. Specifically, this is the one-tailed binomial probability of observing ‘k’ or more successes in ‘n’ trials under H_0:_

The analysis considered the ten possible staircases (2 for each of 5 observers) and found 9 of the 10 staircases followed the predicted pattern (Figure 4). Results showed a remarkably consistent pattern across subjects, pointing toward a significant impact of ordering based on the chromatic background. The chance binomial probability of the results is *p* < 0.001 with n = 10 possible staircases, k = 9 observed staircases, and a chance probability of p = 1/6 of observing the predicted staircase ordering under H_0_.

**Figure 4:**
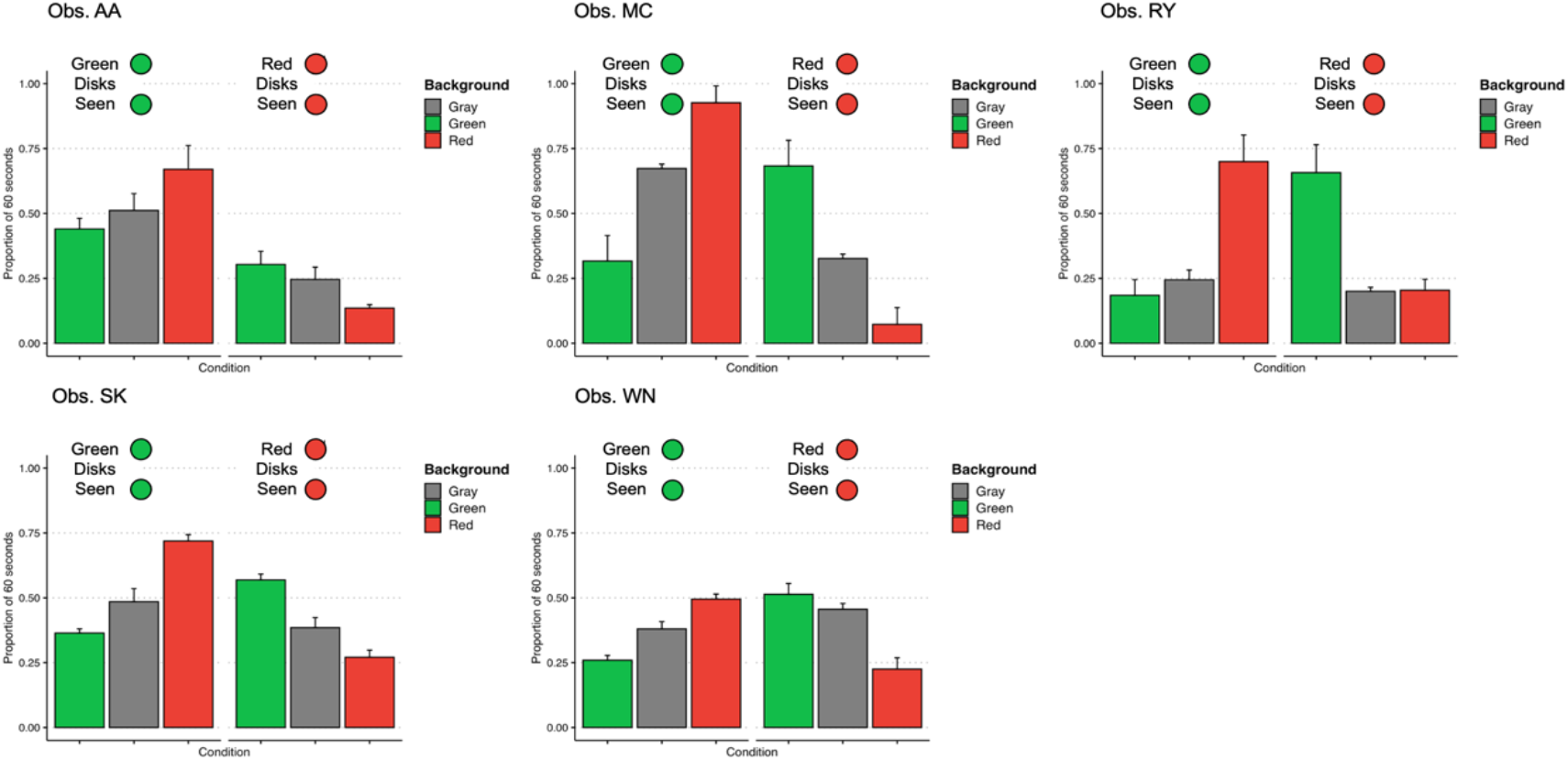
Measurements for five observers. The vertical axis is the proportion of a 60-s trial that each percept was seen. The top horizontal axis groups results by the perceived color. Error bars indicate standard deviation of the mean for measurements taken across three days. Bar color indicates the background color for each measurement.

The results offer clear support for the impact of chromatic background context on perceptual dominance durations, in line with predictions from a chromatically-tuned divisive normalization mechanism. The next experiment considers a potential role of chromatic edge contrast on the observed results.

### Experiment 2

Experiment 2 investigated whether equal-sized neural pools for red and green stimuli would lead to a predominance of similarity-enhanced (grouped) percepts. It was designed to pit a chromatic-contrast-based account against predictions from the tuned-divisive normalization model. Stimulus chromatic contrast was held constant across this and the previous experiment, so if contrast were the primary driver, then rivalrous disks and backgrounds should predominantly resolve as difference-enhanced, with chromatic contrast guiding perceptual selection. On the other hand, pooled-divisive normalization implies equal-sized neural pools for both red and green signals so rivalrous regions should resolve as the same color more frequently than as different colors.

### Stimuli

The stimuli in Experiment 2 were very similar to those in Experiment 1 except that backgrounds that were dichoptically stable in Experiment 1 but made chromatically rivalrous in Experiment 2 (Figure 5A). There were four measured percepts on each trial since the background and disk regions could each resolve as red or green (e.g., Figure 5 B-E). Percepts were categorized as similarity-enhanced if the disks resolved to be the same color as the background (Figure 5 B-C) or difference-enhanced (Figure 5 D-E) if the disks resolved to be a different color than the background.

**Figure 5:**
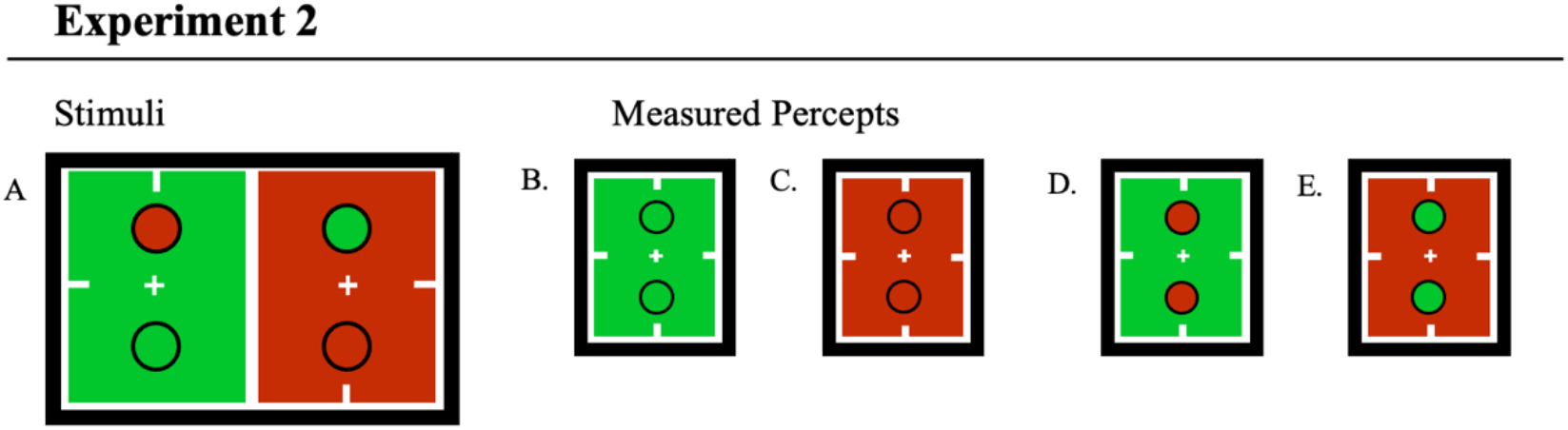
Stimuli and measured percepts for Experiment 2. (A): Rivalrous disks within a rivalrous context. (B-C): Measured similarity-enhanced percepts for stimulus A. (D-E): Measured difference-enhanced percepts for stimulus A.

## Results

Dominance durations were summed separately for similarity-enhanced (Figure 5 B-C) and difference-enhanced (Figure 5 D-E) percepts on each day and then averaged across experimental days. Error bars indicate the standard error across trials. Light [dark] bars represent the average total dominance of difference-enhanced [similarity-enhanced] percepts.

Among the five planned orthogonal contrasts—one for each observer—four were significant (*p* < 0.01) with greater total dominance times for similarity-enhanced percepts (Figure 6). One observer (MC) never reported difference-enhanced percepts, thus displaying a ceiling effect for similarity-enhancement. This observer’s results are consistent with the others’ measurements. In sum, the findings show that neural pools of equal size for red and green signals give predominantly similarity-enhanced percepts, providing further evidence that chromatically tuned divisive normalization mediates perceptual resolution.

**Figure 6:**
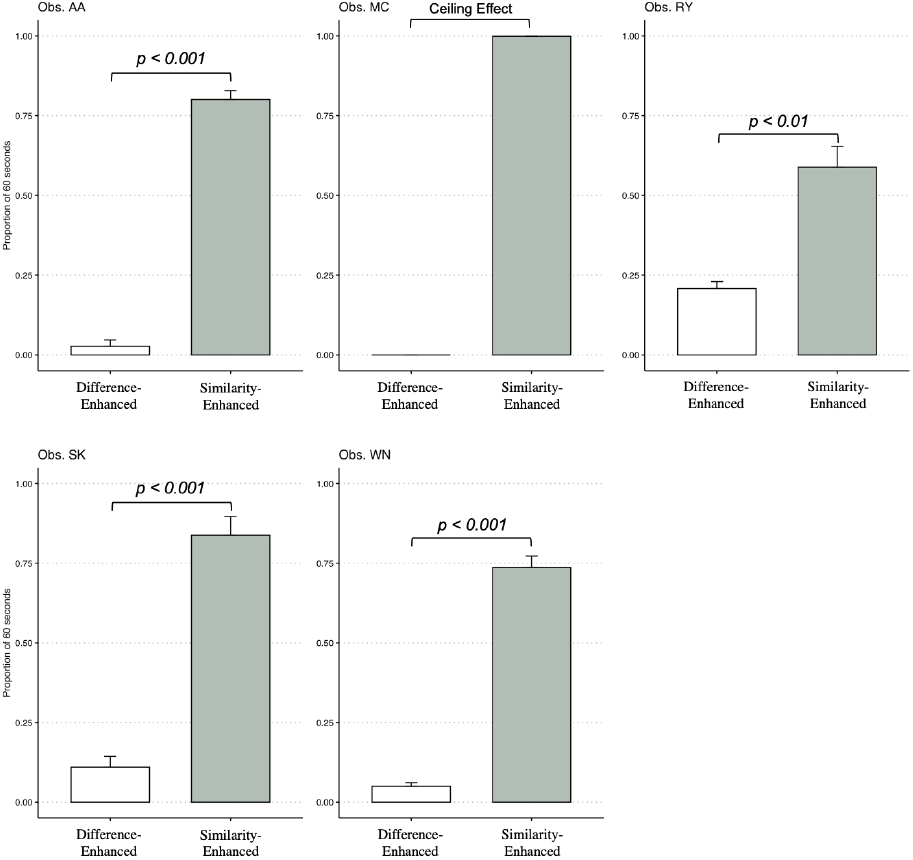
Planned contrasts for five observers. The vertical axis is the proportion of a 60-s trial that each percept was seen. The horizontal axis indicates the response type (“Difference-Enhanced” or “Similarity-Enhanced”). Brackets

## Discussion

This study explored a chromatically tuned divisive normalization account of the perceptual resolution of neural ambiguity during interocular switch rivalry (ISR). The findings provide compelling evidence that the chromatic context of the background influences the dominance durations of percepts, in line with predictions derived from chromatically tuned-divisive normalization. Specifically, the results demonstrate that difference-enhanced percepts are more likely to dominate perception when chromatically contrasted rivalrous regions are presented against stable backgrounds. In contrast, similarity-enhanced percepts are more prevalent when backgrounds are rivalrous. It was hypothesized that difference-enhanced percepts would dominate perception when rivalrous stimuli were presented against a background chromatically congruent with one of the rivalrous chromaticities. This prediction emerges from the proposed framework; specifically, the neural response to the chromatic background is pooled with and attenuates the congruent component of the rivalrous signal. This process, in turn, allows the difference-enhanced representation to dominate perception.

In Experiment 2, similarity-enhanced percepts were hypothesized to dominate perception when both the central disks and the chromatic background were in rivalry. In this case, the two competing neural pools have equal divisive strength, as both include the center and background stimulus regions. Importantly, perceptual resolution in Experiment 2 provides evidence that it is not simple chromatic contrast that drives perceptual resolution of neural ambiguity to be difference-enhanced, as the chromatic contrast is equivalent across both experiments. These findings offer insights into the neural mechanisms underlying interocular grouping and segmentation and suggest potential avenues for further exploration of the interplay between stimulus-driven factors in perceptual selection, such as the spatial distribution of features in a visual stimulus or top-down processes including selective spatial attention.

### Comparison with Similar Models

The results here expand previous research exploring the role of divisive normalization in standard binocular rivalry (Carandini & Heeger, 2012; Moreno-Bote, Rinzel, & Rubin, 2007), selective attention (Lee & Maunsell, 2009; Reynolds & Heeger, 2009), and biased competition models of perception (Desimone & Duncan, 1995; Beck & Kastner, 2009; Peron et al. 2020). These findings also build upon and provide a theoretical foundation for previous work (Peiso & Shevell, 2020). Despite positing divisive normalization as a mechanism supporting biased competition, existing models of binocular rivalry fail to account for all experimental findings. In these cases, divisive normalization provides a mechanism for mutual inhibition by pooling the responses of spatially aligned cells with different feature tuning (Freeman, 2005; Said & Heeger, 2013; Wilson, 2003), but cannot account for interocular grouping and difference enhancement. Moreover, the findings here suggest that antagonistic perceptually selective processes, such as grouping and segmentation, may be supported by a common mechanism, tuned-divisive normalization. This observation warrants further investigation into how the interplay between these processes contributes to the dynamic and flexible nature of visual perception, particularly in complex or ambiguous visual scenes (Palmer & Rock, 1994; Wagemans et al., 2012; Turoman et al., 2021).

A distinguishing feature of the model presented here is that neural representations are pooled based on their response similarities. However, these similarity-linked representations do not universally lead to similarity-enhancement (grouping), as previously predicted. Instead, cells with similar stimulus responses are implicitly pooled together owing to their structural connectivity and current coactivity (Coen-Cagli, Dayan, & Schwartz, 2012; Coen-Cagli, Kohn, & Schwartz, 2015; Shevell & Kingdom, 2008). This structural connectivity might arise from Hebbian plasticity, simply ‘cells that wire together, fire together.’ However, such a rule is implausible in its purest form as it would result in biologically unbounded increases in network activity (Westrick, Heeger, & Landy, 2016). To rectify this, divisive normalization, in conjunction with multiplexed tuning of cells (Akam & Kullmann, 2014), may offer the necessary constraints to a classic Hebbian learning rule, aligning it with observed neural phenomena (Carandini & Heeger, 2012). This framework aligns conceptually with Barlow’s 1981 ‘linking features hypothesis,’ where features are connected by similarity and form non-topographical feature maps. In this model, cell responses implicitly are pooled based on their historical and current activity and mutually exert divisive normalization on each other, thereby reducing the overall response proportional to the size of the divisive pool (Schwartz & Coen-Cagli, 2013). Consequently, the resultant feature maps could facilitate early perceptual organizational processes like figure-ground segregation and exhibit neural nonlinearities such as surround suppression, all through the initial reweighting of feature maps using pooled divisive normalization (Coen-Cagli, Dayan, & Schwartz, 2012; Coen-Cagli, Kohn, & Schwartz, 2015). Similarly, these results and the present framework align with saliency map representations, which capture the relative importance of different visual features in a scene, guiding the allocation of attentional resources (Itti & Koch, 2001). They also are consistent with studies positing that tuned-divisive normalization underlies saliency computations with luminance-defined stimuli (Coen-Cagli, Dayan, & Schwartz, 2012; Schwartz & Simoncelli, 2001; Reynolds & Heeger, 2009). The proposition of early reweighting of the visual signal aligns closest with divisive-normalization accounts of early contextual modulation of perceptual signals (e.g., Schwartz & Simoncelli, 2001; Coen-Cagli, Kohn, & Schwartz, 2015). The overlap with the mechanism described here lends support to the claim that binocular rivalry induces uncommon perceptual experiences via common mechanisms.

The present model diverges from some existing views by negating the need for top-down mechanisms, such as selective attention, to explain interocular grouping (Lee & Blake, 2004; Said & Heeger, 2013). Instead, the model relies on a stimulus-driven, gain-modulatory effect underpinned by feature-tuned divisive normalization that occurs before neural competition between cell populations tuned to incompatible features at the same spatial location. At the same time, attention’s role in perceptual selection cannot be ignored. Continuous sampling by attention drives rivalry dynamics through the adaptation of the dominant neural representation, effectively preventing deterministic outputs. This is consistent with recent evidence suggesting a sampling rate of 4-8 Hz, with higher sampling for single targets than dual targets (Re, Inbar, Richter, & Landau, 2019).

The framework offered here suggests that the mechanisms underlying the resolution of neural ambiguity in binocular rivalry may not be specialized; rather, they result from common processes that reduce redundancy in the visual signal (Coen-Cagli, Dayan, & Schwartz, 2012; Schwartz & Simoncelli, 2001; Solomon, Peirce, & Lennie, 2004). The proposed framework clearly explains our previous finding of difference enhancement in some experimental conditions (Peiso & Shevell, 2020). Consistent with the idea that the visual system employs universal mechanisms to resolve unique instances of neural ambiguity, the divisive normalization model postulates that responses from similarly tuned cells are pooled across the visual field. Following this initial reweighting of the visual signal, the processed information advances to a second stage, where the dominant representation attains visual awareness. A mechanism rooted in spatially tuned normalization, like spatial attention, may be essential for influencing neural responses from neurons with divergent tuning characteristics, and therefore lack the structural connectivity to support implicit response pooling. The attentional mechanism then inherits re-weighted signals from the initial processing stage, ultimately leading to the biased selection that shapes perception. The two-factor model proposed by Brascamp et al., (2019) offers a descriptive account of interocular grouping as an outcome of domain-general spatial integration. Our model posits that interocular grouping would occur earlier in the processing stream via feature-tuned normalization pools prior to neural competition related to interocular conflict.

These results offer insights into the role of tuned-divisive normalization in the perceptual resolution of neural ambiguity, some limitations warrant mention. First, the experiments primarily focused on chromatic context, which may not fully capture the complexities of other visual features, such as luminance or motion. Second, the model, though predictive, is based on a simplified representation of neural interactions and may not account for more complex neural dynamics. Nevertheless, the measurements here are consistent with a tuned-divisive normalization mechanism that supports the brain’s capacity for employing contextual information when resolving neural ambiguity. The findings can inform the development of computational models aimed at understanding how early perceptual organization creates biases in what we see.

## Acknowledgements

Linda Glennie

Sunny Lee (Data Collection)

Research reported in this publication was supported by National Eye Institute of the National Institutes of Health under award number R01EY026618 (SS).

This material is based upon work supported by the National Science Foundation under Grant No. 1746045 (JP)

This work was supported by the National Science Foundation through the Physics Frontier Center for Living Systems PHY-2317138 (SEP).

## References

1. Brascamp, J. W., & Shevell, S. K. (2021). The certainty of ambiguity in visual neural representations. Annual Review of Vision Science, 7, 465–486.

2. Coen-Cagli, R., Kohn, A., & Schwartz, O. (2015). Flexible gating of contextual influences in natural vision. Nature neuroscience, 18(11), 1648–1655.

3. Lee, T. S., & Mumford, D. (2003). Hierarchical Bayesian inference in the visual cortex. JOSA A, 20(7), 1434–1448.

4. Fowlkes, C. C., Martin, D. R., & Malik, J. (2007). Local figure–ground cues are valid for natural images. Journal of vision, 7(8), 2–2.

5. Desimone, R., & Duncan, J. (1995). Neural mechanisms of selective visual attention. Annual review of neuroscience, 18(1), 193–222.

6. Hillyard, S. A., Vogel, E. K., & Luck, S. J. (1998). Sensory gain control (amplification) as a mechanism of selective attention: electrophysiological and neuroimaging evidence. Philosophical Transactions of the Royal Society of London. Series B: Biological Sciences, 353(1373), 1257–1270.

7. Ling, S., & Blake, R. (2012). Normalization regulates competition for visual awareness. Neuron, 75(3), 531–540.

8. Murray, R. F., & Adams, W. J. (2019). Visual perception and natural illumination. Current Opinion in Behavioral Sciences, 30, 48–54.

9. Brascamp, J. W., Qian, C. S., Hambrick, D. Z., & Becker, M. W. (2019). Individual differences point to two separate processes involved in the resolution of binocular rivalry. Journal of Vision, 19(12), 15–15.

10. Cao, T., Wang, L., Sun, Z., Engel, S. A., & He, S. (2018). The independent and shared mechanisms of intrinsic brain dynamics: Insights from bistable perception. Frontiers in Psychology, 9, 589.

11. Buckthought, A., Kirsch, L. E., Fesi, J. D., & Mendola, J. D. (2021). Interocular grouping in perceptual rivalry localized with fMRI. Brain Topography, 34, 323–336.

12. Lee, S. H., & Blake, R. (1999). Rival ideas about binocular rivalry. Vision research, 39(8), 1447–1454.

13. Lee, S. H., & Blake, R. (2004). A fresh look at interocular grouping during binocular rivalry. Vision research, 44(10), 983–991.

14. Peiso, J. R., & Shevell, S. K. (2020). Seeing fruit on trees: enhanced perceptual dissimilarity from multiple ambiguous neural representations. JOSA A, 37(4), A255–A261.

15. Louie, K., & Glimcher, P. W. (2019). Normalization principles in computational neuroscience. In Oxford Research Encyclopedia of Neuroscience.

16. Schwartz, O., & Simoncelli, E. P. (2001). Natural signal statistics and sensory gain control. Nature neuroscience, 4(8), 819–825.

17. Carandini, M., & Heeger, D. J. (2012). Normalization as a canonical neural computation. Nature Reviews Neuroscience, 13(1), 51–62.

18. Olsen, S. R., Bhandawat, V., & Wilson, R. I. (2010). Divisive normalization in olfactory population codes. Neuron, 66(2), 287–299.

19. Schwartz, O., & Simoncelli, E. (2000). Natural sound statistics and divisive normalization in the auditory system. Advances in neural information processing systems, 13.

20. Heeger, D. J. (1992). Normalization of cell responses in cat striate cortex. Visual neuroscience, 9(2), 181–197.

21. Carandini, M., Heeger, D. J., & Movshon, J. A. (1997). Linearity and normalization in simple cells of the macaque primary visual cortex. Journal of Neuroscience, 17(21), 8621–8644.

22. Barlow, H. B. (1961). Possible principles underlying the transformation of sensory messages. Sensory communication, 1(01), 217–233.

23. Simoncelli, E. P., & Schwartz, O. (1999). Modeling surround suppression in V1 neurons with a statistically derived normalization model. Advances in neural information processing systems, 153–159.

24. Brouwer, G. J., & Heeger, D. J. (2011). Cross-orientation suppression in human visual cortex. Journal of neurophysiology, 106(5), 2108–2119.

25. Freeman, A. W. (2005). Multistage model for binocular rivalry. Journal of neurophysiology, 94(6), 4412–4420.

26. Lehky, S. R. (1988). An astable multivibrator model of binocular rivalry. Perception, 17(2), 215–228.

27. Wilson, H. R. (2003). Computational evidence for a rivalry hierarchy in vision. Proceedings of the National Academy of Sciences, 100(24), 14499–14503.

28. Said, C. P., & Heeger, D. J. (2013). A model of binocular rivalry and cross-orientation suppression. PLoS computational biology, 9(3), e1002991.

29. Bloem, I. M., & Ling, S. (2019). Normalization governs attentional modulation within human visual cortex. Nature communications, 10(1), 5660.

30. Klímová, M., Bloem, I. M., & Ling, S. (2023). Attention preserves the selectivity of feature-tuned normalization. Journal of Neurophysiology, 130(4), 990–998.

31. Slezak, E., & Shevell, S. K. (2018). Perceptual resolution of color for multiple chromatically ambiguous objects. JOSA A, 35(4), B85–B91.

32. Brascamp, J. W., Van Ee, R., Noest, A. J., Jacobs, R. H., & van den Berg, A. V. (2006). The time course of binocular rivalry reveals a fundamental role of noise. Journal of vision, 6(11), 8–8.

33. Kim, Y. J., Grabowecky, M., & Suzuki, S. (2006). Stochastic resonance in binocular rivalry. Vision research, 46(3), 392–406.

34. Moreno-Bote, R., Rinzel, J., & Rubin, N. (2007). Noise-induced alternations in an attractor network model of perceptual bistability. Journal of neurophysiology, 98(3), 1125–1139.

35. Lankheet, M. J. (2006). Unraveling adaptation and mutual inhibition in perceptual rivalry. Journal of Vision, 6(4), 1–1.

36. van Ee, R. (2009). Stochastic variations in sensory awareness are driven by noisy neuronal adaptation: evidence from serial correlations in perceptual bistability. JOSA A, 26(12), 2612–2622.

37. Wilson, H. R., Blake, R., & Lee, S. H. (2001). Dynamics of travelling waves in visual perception. Nature, 412(6850), 907–910.

38. Brascamp, J. W., & Blake, R. (2012). Inattention abolishes binocular rivalry: Perceptual evidence. Psychological Science, 23(10), 1159–1167.

39. Dieter, K. C., Brascamp, J., Tadin, D., & Blake, R. (2016). Does visual attention drive the dynamics of bistable perception?. Attention, Perception, & Psychophysics, 78, 1861–1873.

40. Hancock, S., & Andrews, T. J. (2007). The role of voluntary and involuntary attention in selecting perceptual dominance during binocular rivalry. Perception, 36(2), 288–298

41. Li, H. H., Rankin, J., Rinzel, J., Carrasco, M., & Heeger, D. J. (2017). Attention model of binocular rivalry. Proceedings of the National Academy of Sciences, 114(30), E6192–E6201.

42. Coen-Cagli, R., Dayan, P., & Schwartz, O. (2012). Cortical surround interactions and perceptual salience via natural scene statistics. PLoS computational biology, 8(3), e1002405.

43. Shevell, S. K., & Kingdom, F. A. (2008). Color in complex scenes. Annu. Rev. Psychol., 59, 143–166. Kovács, I., Papathomas, T. V., Yang, M., & Fehér, Á. (1996). When the brain changes its mind: Interocular grouping during binocular rivalry. Proceedings of the National Academy of Sciences, 93(26), 15508–15511.

44. Westrick, Z. M., Heeger, D. J., & Landy, M. S. (2016). Pattern adaptation and normalization reweighting. Journal of Neuroscience, 36(38), 9805–9816.

45. Akam, T., & Kullmann, D. M. (2014). Oscillatory multiplexing of population codes for selective communication in the mammalian brain. Nature Reviews Neuroscience, 15(2), 111–122.

46. Barlow, H. B. (1981). The ferrier lecture, 1980. Proceedings of the Royal Society of London. Series B. Biological Sciences, 212(1186), 1–34.

47. Re, D., Inbar, M., Richter, C. G., & Landau, A. N. (2019). Feature-based attention samples stimuli rhythmically. Current Biology, 29(4), 693–699.

48. Solomon, S. G., Peirce, J. W., & Lennie, P. (2004). The impact of suppressive surrounds on chromatic properties of cortical neurons. Journal of Neuroscience, 24(1), 148–160.

49. Ni, A. M., & Maunsell, J. H. (2019). Neuronal effects of spatial and feature attention differ due to normalization. Journal of Neuroscience, 39(28), 5493–5505.

50. Reynolds, J. H., & Heeger, D. J. (2009). The normalization model of attention. Neuron, 61(2), 168–185.

51. Moradi, F., & Heeger, D. J. (2009). Inter-ocular contrast normalization in human visual cortex. Journal of vision, 9(3), 13–13.

52. Lee, B. B., Martin, P. R., & Valberg, A. (1988). The physiological basis of heterochromatic flicker photometry demonstrated in the ganglion cells of the macaque retina. The Journal of physiology, 404(1), 323–347.

53. Bone, R. A., & Landrum, J. T. (2004). Heterochromatic flicker photometry. Archives of biochemistry and biophysics, 430(2), 137–142.

54. Wyszecki, G. & Stiles, W. S. (1982). Color scienceIconcepts and methods, quantitative data and formulae, 2nd edn. New York: John Wiley.

55. Christiansen, J. H., D’Antona, A. D., & Shevell, S. K. (2017). Chromatic interocular-switch rivalry. Journal of vision, 17(5), 9–9.

56. Logothetis, N. K., Leopold, D. A., & Sheinberg, D. L. (1996). What is rivalling during binocular rivalry?. Nature, 380(6575), 621–624.

57. Lange, R., & Shevell, S. K. (2020). Does feature integration affect resolution of multiple simultaneous forms of ambiguity?. JOSA A, 37(4), A105–A113.

58. MacLeod, D. I., & Boynton, R. M. (1979). Chromaticity diagram showing cone excitation by stimuli of equal luminance. JOSA, 69(8), 1183–1186.

59. Carter, O., & Cavanagh, P. (2007). Onset rivalry: Brief presentation isolates an early independent phase of perceptual competition. PloS one, 2(4), e343.

60. Lee, J., & Maunsell, J. H. (2009). A normalization model of attentional modulation of single unit responses. PloS one, 4(2), e4651.

61. Beck, D. M., & Kastner, S. (2009). Top-down and bottom-up mechanisms in biasing competition in the human brain. Vision research, 49(10), 1154–1165.

62. Peron, S., Pancholi, R., Voelcker, B., Wittenbach, J. D., Ólafsdóttir, H. F., Freeman, J., & Svoboda, K. (2020). Recurrent interactions in local cortical circuits. Nature, 579(7798), 256–259.

63. Palmer, S., & Rock, I. (1994). Rethinking perceptual organization: The role of uniform connectedness. Psychonomic bulletin & review, 1(1), 29–55.

64. Wagemans, J., Elder, J. H., Kubovy, M., Palmer, S. E., Peterson, M. A., Singh, M., & von der Heydt, R. (2012). A century of Gestalt psychology in visual perception: I. Perceptual grouping and figure– ground organization. Psychological bulletin, 138(6), 1172.

65. Turoman, N., Tivadar, R. I., Retsa, C., Murray, M. M., & Matusz, P. J. (2021). Towards understanding how we pay attention in naturalistic visual search settings. NeuroImage, 244, 118556.

66. Itti, L., & Koch, C. (2001). Feature combination strategies for saliency-based visual attention systems. Journal of Electronic imaging, 10(1), 161–169.

67. Lee, J., & Maunsell, J. H. (2009). A normalization model of attentional modulation of single unit responses. PloS one, 4(2), e4651.

